# Critical Assessment of ML models for ADMET Prediction in TDC leaderboards

**DOI:** 10.64898/2026.02.26.708193

**Authors:** Ihor Koleiev, Roman Stratiichuk, Nazar Shevchuk, Mykola Melnychenko, Oleksiy Nyporko, Daniil Todoryshyn, Vladyslav Husak, Sergii Starosyla, Semen Yesylevskyy, Alan Nafiiev

## Abstract

In this work we performed a critical assessment of the benchmarking procedures used in Therapeutics Data Commons (TDC) ADMET leaderboards, focusing on reproducibility, robustness against data leakage, and signs of test-set overfitting across all 22 TDC ADMET endpoints. For each endpoint, the top 3 leaderboard models were screened with a unified protocol: execution environment reproducibility check, data leakage assessment, verification of hyperparameter optimisation practices, and final re-evaluation of results and TDC ranking. Only 3 methods (CaliciBoost, MapLight, MapLight+GNN) passed all checks and showed overall reproducible performance, whereas most of top-ranked models exhibited unavailable code, non-reproducible execution environments, runtime incompatibilities, or various methodological flaws. In particular, we identified direct or indirect data leakages in MiniMol, GradientBoost and XGBoost models. We also used our in-house models based on the Mol2Vec architecture to investigate the consequences of deliberately overfitting on the TDC test set. It is shown that deliberate or accidental tuning on the public test set may lead to significant inflation of the model metrics and leaderboard position. Our results emphasize the urgent need for better public ADMET benchmarks with the hidden test sets, strict dataset versioning and model submission with standardized inference environments.

## Introduction

The process of novel drug development is lengthy, costly, and associated with a high risk of failure at late stages (1,2). One of the primary reasons for attrition (accounting for up to 50% of failures) is not insufficient therapeutic efficacy, but rather unfavorable pharmacokinetic and toxicological properties, collectively referred to as ADMET (absorption, distribution, metabolism, excretion, and toxicity) (3–5).

Understanding the ADMET profile at early stages of drug development is critically important for rational drug design, as it enables the elimination of inherently non-viable compounds before performing expensive *in vivo* testing (6). Preliminary evaluation of chemical libraries allows researchers to focus on regions of chemical space that exhibit an optimal balance between biological activity and safety (7,8), thereby dramatically reducing both drug development time and costs.

Historically, several computational approaches have been developed for ADMET assessment, ranging from classical “drug-likeness” rules (9) to quantitative structure–property relationship (QSAR) methods based on linear regression. However, with the accumulation of large-scale data in public databases (such as ChEMBL, PubChem, and ZINC) and the increasing complexity of prediction tasks, the focus has shifted toward machine learning (ML) methods. Today, ML models are capturing an ever-growing segment of the drug discovery landscape, gradually displacing traditional physicochemical simulations (10). This trend is driven by their ability to capture nonlinear relationships in high-dimensional data and their high prediction throughput, which is suitable for large-scale screenings (11–13). The most recent ML ADMET prediction models exploit novel neural network architectures such as graph neural networks (GNNs) (14,15) and transformers (11).

With rapid proliferation of ADMET prediction algorithms, an urgent need has emerged for rigorous and fair comparison of their quality (16,17). Traditional evaluation strategies include cross-validation and testing on external datasets using standard performance metrics (RMSE, MAE for regression; AUC-ROC, F1-score for classification) (18). While these approaches are implemented in widely used libraries such as scikit-learn and DeepChem, their application to chemical data requires careful consideration of the features of chemical space structure: models often demonstrate excellent performance on familiar molecular scaffolds but lose predictive power when confronted with fundamentally novel chemotypes (19–21).

Dedicated online ADMET benchmarking platforms have gained significant traction in recent years. Notable examples include MoleculeNet (22), PharmaBench (23), and arguably the most well-known Therapeutic Data Commons (TDC) (24). The TDC platform is a comprehensive resource that provides access to curated ADMET datasets with pre-defined data splits and introduces a public leaderboard for ADMET prediction models. As an open data platform, TDC is often referred to as the ‘gold standard’ for the academic community, allowing researchers to instantly compare their results with the cutting-edge rivals (25,26).

However, the availability of fully open datasets is both an advantage and a serious drawback. Open train-test split introduces the risk of deliberate cheating or unintentional model adaptation to a specific test set (27,28). When a test set remains publicly available for years, researchers may implicitly optimize architectures or hyperparameters to maximize performance on that specific dataset, without guaranteeing robustness under real-world conditions involving truly novel data (29). Moreover, some recent ML models reach prediction accuracy for certain endpoints that exceeds the accuracy of the experimental assays used to generate the training data. This phenomenon is a clear indicator of overfitting: the model captures dataset-specific noise or systematic measurement errors rather than genuine biological or chemical relationships (30,31).

Researchers may resort to intentional overfitting because they need to demonstrate state-of-the-art results to get their articles accepted by high-ranking journals, obtain research grants, boost their own careers or promote commercialisation of the models in the drug discovery market. Regardless of the reason, inflated leaderboard positions diminish scientific value of the benchmarks and hamper the progress in the field.

To flag overfitting, it is important to understand what the maximum ‘fair’ accuracy of predicting ADMET properties is. Given experimental uncertainty and inter-laboratory variability (32), the coefficient of determination R^2^ for most biological endpoints rarely exceeds 0.7–0.8 (33). As emphasized by domain experts, any model claiming accuracy beyond this threshold on noisy biological data is likely learning random fluctuations in a particular dataset rather than fundamental chemical principles (34,35).

Thus, at the current stage of cheminformatics development, there is an urgent need for a critical assessment of existing ADMET benchmarking practices. The field must transition from merely recording leaderboard rankings toward the development of robust evaluation methodologies that explicitly account for data uncertainty (16,36) and prevent deliberate or unintentional data manipulations.

The objective of the present study is to perform a critical analysis of the benchmarking procedures provided by the TDC platform. We performed assessment of the top TDC leaderboard models for all TDC deposed ADMET endpoints paying special attention to the following aspects:

- Reproducibility of the model training and inference.
- Robustness of the model training protocol in avoiding data leakage between test and train sets.
- Signs of overfitting to test or otherwise inflating the metrics.

We show that multiple top-ranked models in TDC leaderboards have serious issues in one or several of these categories and thus could not be considered as the state-of-the-art in the field despite seemingly outstanding performance metrics.

As an example of a properly developed and fairly tested ML model for ADMET prediction we also provide the results of our in-house models based on Mol2Vec architecture that feature decent positions in TDC leaderboards.

## Materials and Methods

Among the ADMET property prediction models displayed on the TDC leaderboards, top models were identified for each of the 22 ADMET endpoints presented here. After comparing the lists of top models and identifying/removing duplicates, a final list of models was formed for subsequent quality assessment. The models were checked according to a procedure including verifying model reproducibility, assessing potential data leakage, and checking the correctness of model optimization. At each stage, models that did not meet the relevant criteria (see below) were eliminated, saving resources and evaluation time. Only models that successfully passed all stages of preliminary testing were admitted to evaluation using the complete set of ADMET endpoints. The ML model we developed for evaluating ADMET properties was also tested on this same complete set.

### 1. Analysis of ADMET Benchmarks

#### 1.1. Selection of leaderboard models

For each ADMET endpoint, we focused on the top three leaderboard models because benchmark analyses typically concentrate on the small group of leading methods rather than the full ranking. In many TDC endpoints, performance differences among the top models are small, so restricting the analysis to the top three captures the main competitive approaches while avoiding over-representation of closely related variants. This choice therefore allows the study to remain both representative of state-of-the-art performance and practically feasible for detailed verification. After removing duplicates across endpoints, this procedure resulted in a final set of ten representative architectures:

ADMETrix, CFA, GradientBoost+, MapLight, Maplight + GNN, MiniMol, CaliciBoost, SimGCN, XGBoost, and ZairaChem.

This strategy ensured that the analysis focused on models that are competitive according to the community of TCD benchmarks and, simultaneously, lets rational use the resource and time for obtaining maximum useful information.

#### 1.2. ADMET endpoints

To evaluate the models, all 22 endpoints available within the TDC ADMET benchmark leaderboard were involved. These endpoints span the full ADMET spectrum - absorption, distribution, metabolism, excretion, and toxicity - and include both regression and classification tasks with task-specific evaluation metrics (MAE, Spearman correlation, AUROC, AUPRC). A complete list of the endpoints, dataset sizes, and metrics is provided in Supplementary materials (Table S1).

For individual models, evaluation was stopped early if significant issues (e.g. impossibility to code running, suggested data leakage or unclear procedure of hyperparameter tuning) were encountered at earlier validation stages.

#### 1.3. Dataset sources and splits

All datasets were obtained directly from TDC and used with the official training/test splits provided by the TDC platform. No resplitting, rebalancing, or data augmentation was applied, ensuring comparability with leaderboard results.

An exception was the MiniMol model, which is a foundation model pretrained on the LargeMix dataset. In this case, the LargeMix pretraining set was filtered according to the developers’ recommendations by removing all molecules present in the TDC test sets, thereby preventing direct information leakage from pretraining to evaluation.

### 2. Evaluation Protocol for Leaderboard Models

All selected leaderboard models were assessed using a unified, multi-stage protocol designed to ensure reproducibility, methodological correctness, and fair comparison.

#### 2.1. Reproducibility check

Models were tested in a local environment. Reproducibility was assessed using a single endpoint per model: specifically, the endpoint on which the model achieved its highest leaderboard rank. For this endpoint, the reported leaderboard metric was recomputed and compared with the published value. Given the sensitivity of some models to random seeds and software versions, minor numerical deviations in absolute metric values were considered acceptable, provided that they did not affect the model’s relative ranking on the leaderboard; only systematic discrepancies leading to rank changes were considered as essential.

#### 2.2. Assessment of data leakage

To assess potential data leakage between training and test datasets, Tanimoto similarity scores were computed using Morgan fingerprints (radius 3, 2048 bits). Pairwise similarities were calculated between all molecules in the training and test sets.

Two complementary statistical metrics were analysed:

a) Median similarity, reflecting the overall overlap between chemical spaces.
b) Maximum similarity, highlighting the presence of highly similar or identical molecules (Tanimoto ∼ 1).

For the MiniMol model, similarity analysis was conducted between the pretraining dataset and the union of all TDC test molecules. An additional qualitative check was performed by querying PubChem for known ADMET-related proteins associated with the compounds from the pretraining dataset.

#### 2.3. Verification of optimisation procedures

Primary publications and software repositories were examined to verify that:

- Hyperparameter optimization procedures were explicitly described.
- No tuning or model selection was performed on the test sets.

This step is essential because all TDC test sets are publicly accessible, making intentional or accidental overfitting to a test set possible.

#### 2.4. Final evaluation

Models that passed the previous stages were evaluated following the official TDC instructions (https://tdcommons.ai/benchmark/overview/).

The resulting metrics were compared against all leaderboard entries available at the time of writing (November 2025), and model rankings were recomputed accordingly.

### 3. Development of an In-House ML models

To complement the analysis of leaderboard models, the in-house ML models were developed and evaluated using the TDC protocol.

#### 3.1. Model architecture

LightGBM (Light Gradient Boosting Machine) was selected as the base learning algorithm due to its computational efficiency, efficient handling of sparse molecular fingerprints, and consistently strong baseline performance in QSAR and ADMET prediction tasks. LightGBM implements gradient boosting decision trees with histogram-based split finding, which enables efficient training even for high-dimensional and heterogeneous molecular feature spaces. Regression tasks were handled using LGBMRegressor (37) API, while classification tasks employed LGBMClassifier (38) API with a binary cross-entropy objective.

#### 3.2. Molecular representations

Mol2Vec embeddings were used as a fixed baseline representation across all benchmarks. Mol2Vec is an unsupervised embedding method inspired by distributional semantics models in natural language processing. In this approach, molecular substructures are treated analogously to words, while molecules are represented as sentences of such substructures. Embeddings are learned such that substructures occurring in similar chemical contexts are mapped to nearby points in a continuous vector space.

Mol2Vec embeddings were generated using a Skip-gram Word2Vec model trained on a large and chemically diverse set of 883,897,271 small molecules. Substructures were identified using Morgan fingerprints with radius 0 and 1, and the resulting vocabulary encompassed 13,278 unique “words.” Each molecule was represented by aggregating the embeddings of its constituent substructures into a single 512-dimensional vector. Substructures not present in the training vocabulary were assigned a default embedding corresponding to the mean vector of the embedding space.

In addition to Mol2Vec, diverse sets of molecular fingerprints (23 types, table S2) were used as complementary feature groups to define the optimal feature set for calculation of each endpoint. In addition, benchmark-specific sets of features “filtered_descs” were generated by combining RDKit 2D and Mordred 2D descriptors with subsequent filtering. The filtering procedure included removing duplicate descriptors, excluding descriptors with calculation errors, low-variance filtering, and rejecting features with high correlation (cross-correlation filtering). Descriptor filtering was performed using training data only to avoid information leakage.

#### 3.3. Optimization pipeline

Model optimization followed a two-phase procedure. In Phase 1, Sequential Forward Selection (SFS) was applied to identify an optimal set of molecular feature groups for each benchmark. Starting from a fixed Mol2Vec baseline, additional feature groups were iteratively added based on five-fold cross-validation performance. The selection criterion was the selection_score value, calculated as the difference between the mean value of the appropriate metric and the corresponding standard deviation.

In Phase 2, Bayesian hyperparameter optimization was performed using the Optuna framework (39) with a Tree-structured Parzen Estimator sampler (40). Hyperparameters were optimised using the same cross-validation protocol as in Phase 1. Optimisation was skipped for benchmarks where the model already achieved top leaderboard rank after feature selection alone.

Performance on a test set was monitored exclusively for verification purposes and was never used for feature selection or hyperparameter tuning.

#### 3.4. Final training and evaluation

For each benchmark, the final model was re-trained on the full training set using the selected features and optimised hyperparameters, and then evaluated on the TDC test set. To explicitly quantify the contribution of successive optimisation stages, all 22 ADMET endpoints were evaluated three times for the in-house models: (i) using the baseline (non-optimised) model, (ii) after the feature selection stage (Phase 1 optimisation), and (iii) after hyperparameter optimisation (Phase 2 optimisation).

#### 3.5 Development of deliberately overfitted variants of In-house ML models

To evaluate the possible overfit influence on the performance metrics and appropriate model ranks in TDC ADMET leaderboards, deliberately overfitted variants of In-house ML models were constructed. Overfitting was introduced by using the TDC test datasets instead of TDC train datasets during the both SFS and HPO optimisation phases. Resulting overfitted models have features sets and hyperparameters different from honest ones (see Supplementary tables S3 and S4).

## Results

### Checking availability and reproducibility of the models

First, we assessed the code availability and possibility to reproduce the results and have found critical issues in a number of models.

Source code of the model **CFA** (Combinatorial Fusion Analysis) model (41) was not accessible due to the broken links on both the TDS website and the corresponding paper. This model ranks first in the *clearance_hepatocyte_az* and *half_life_obach* leaderboards, second in the *bbb_martins* and *herg* leaderboards, and third in the *dili* and *vdss_lombardo* leaderboards.

For **ADMETrix** (42) (first in the *ld50_zhu*, second in the *ames* and *lipophilicity_astrazeneca*, and third in the *cyp2c9_veith*) and **SimGCN** models (43) (third in the *herg*) the execution environments could not be reproduced based on the available installation instructions. In the case of ADMETRix, the provided setup description specifies a broad Python version requirements, whereas the dependency set implicitly assumes a narrower and internally inconsistent range of versions; as a result, several required packages were either incompatible or unavailable, preventing successful environment construction. For SimGCN, the installation instructions do not specify a compatible version-matched combination of PyTorch and PyTorch Geometric extensions. This leads to unresolved dependencies and makes environment construction impossible. Fragments of the logs with the corresponding errors are shown in the Supplementary Information.

**ZairaChem** model (44) (first in the *ames*, second in the *cyp2c9_substrate_carbonmangels* and *dili*, third in the *bioavailability_ma*) was successfully deployed, but execution failed due to incompatibilities among core libraries in the inference pipeline. In particular, the installation procedure delegates environment setup to an external script without fixing compatible versions of PyTorch and the TabPFN components used by the model, which results in runtime import errors despite nominally successful installation. The relevant log fragment containing a description of the errors encountered is provided in the Supplementary Information.

These findings emphasize the lack of proper software development practices, testing and quality control, which is, unfortunately, common in the academic machine learning community and hinders adoption and cross-validation of developed models.

### Assessment of data leakage between training and test datasets

**MiniMol** is one of the most successful models, which is ranked first on seven TDC leaderboard (*bbb_martins, bioavailability_ma, cyp2c9_substrate_carbonmangels, dili, hia_hou, lipophilicity_astrazeneca and solubility_aqsoldb*). MiniMol is a foundational model pretrained on the massive Graphium **LargeMix** dataset containing approximately six million unique molecules. According to the authors, molecules present in all TDC ADMET test sets were removed from the pretraining corpus to prevent data leakage. However, the paper does not specify precisely how this matching and filtering were performed, which raises concerns about potential data leakage

The LargeMix dataset comprises three types of data: transcriptomic data constituting about 5% of the pretraining set (L1000 VCAP and L1000 MCF7), biochemical assay data constituting about 41% (PCBA-1328), and quantum chemical data constituting about **54%** (PCQM4M G25 and PCQM4M N4 datasets).

Following the description in the original paper, canonical SMILES were used to remove molecules belonging to the TDC test sets from the MiniMol pretraining corpus. This results in exclusion rates consistent with those reported by the MiniMol developers — 7% for MCF7, 4% for VCAP, and 0.6% for PCBA. However, inspection of similarity distributions (Figs. 1A-C) demonstrates the presence of molecules that are highly similar or identical to ones in the TDC test set across all three analyzed MiniMol training datasets with the largest number of such molecules observed in the PCBA dataset.

**Figure 1.**
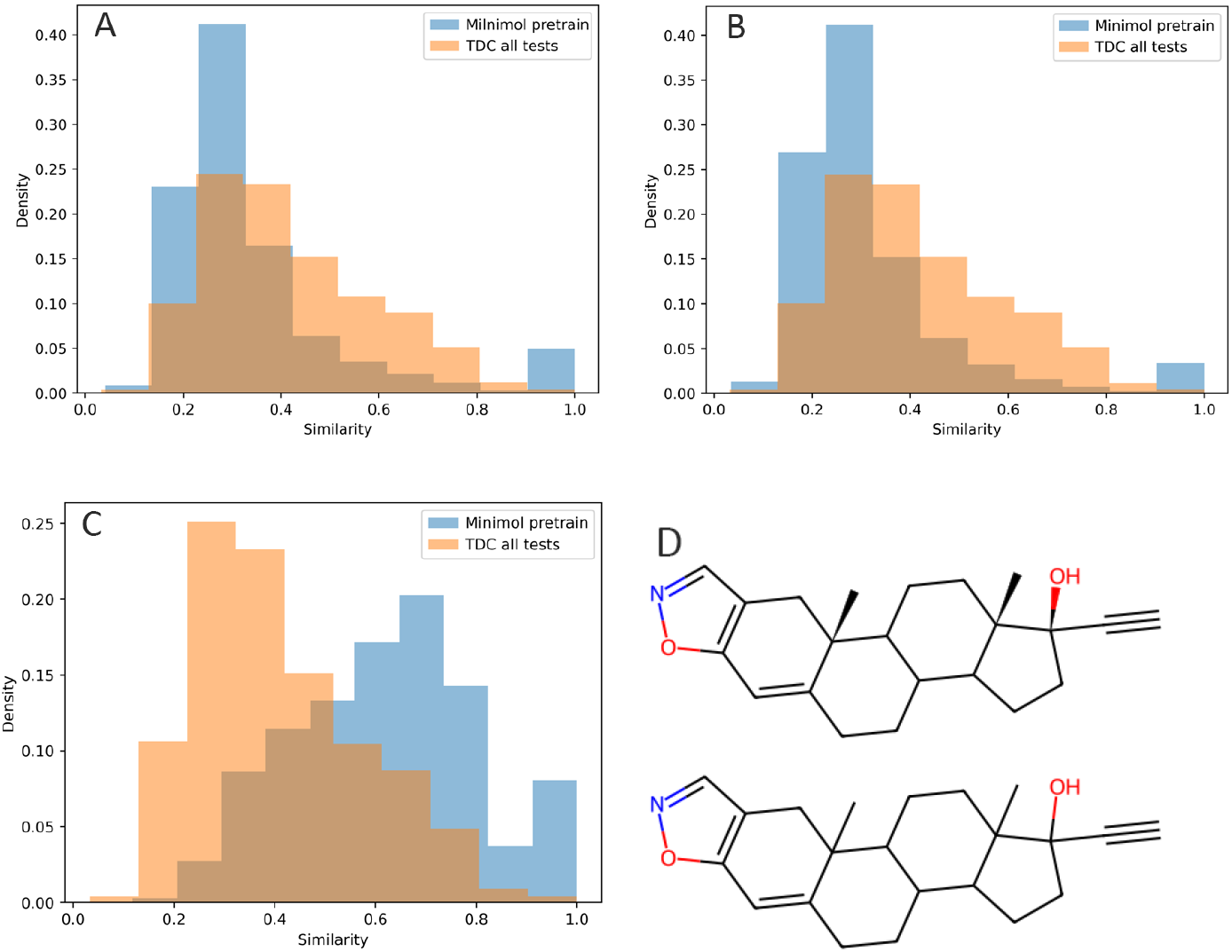
Distribution of similarities between Minimol pretrain and TDC test sets. (A) - MCF7, (B) - VCAP, (C) - PCBA. (D) - danazol molecule as found in TDC all-test dataset (with chirality annotations, top) and MiniMol pretrain dataset (without chirality annotations, bottom).

Upon closer inspection it became evident that simple filtering by canonical SMILES is insufficient, as it often fails to account for molecular chirality properly. It removes only one specific stereoisomer from the dataset, while alternative stereoisomers and/or representations of the same compound with missing stereochemistry annotations are retained and lead to a data leakage. For example, danazol is present as a concrete stereoisomer in the TDC all-test dataset (Figure 2a), yet it was not removed from the MiniMol pretraining dataset during filtering because its stereochemistry was not explicitly specified in the MiniMol training set (Figure 1D). Tautomerism may also contribute to the persistence of undetected duplicates, as different tautomeric forms of the same compound are usually represented by distinct canonical SMILES strings. Consequently, molecules that are chemically equivalent but differ only in tautomeric state may bypass filtering and contribute to unintended data leakage.

### Verification of hyperparameter optimisation procedures

At this validation stage, significant issues were detected in two models — **GradientBoost** and **XGBoost**, both developed by the same author. GradientBoost ranks first on the *ppbr_az* leaderboard, whereas XGBoost ranks second on the *caco2_wang* leaderboard.

For both models we identified a flaw in the dataset splitting procedure applied during hyperparameter optimization: the validation sets generated at the model tuning stage partially overlapped with the (public) TDC test sets. This issue originates from an incorrect function call that, prior to hyperparameter optimization, re-splits the full dataset randomly into training, validation, and test subsets with the default hard-coded random seed of 42. As a consequence, the validation set contains, among others, molecules from the original TDC test set constituting a direct data leakage. The similarity distributions between the test set and the validation sets used by the model authors and corrected by us are shown in Fig. 3.

**Figure 3.**
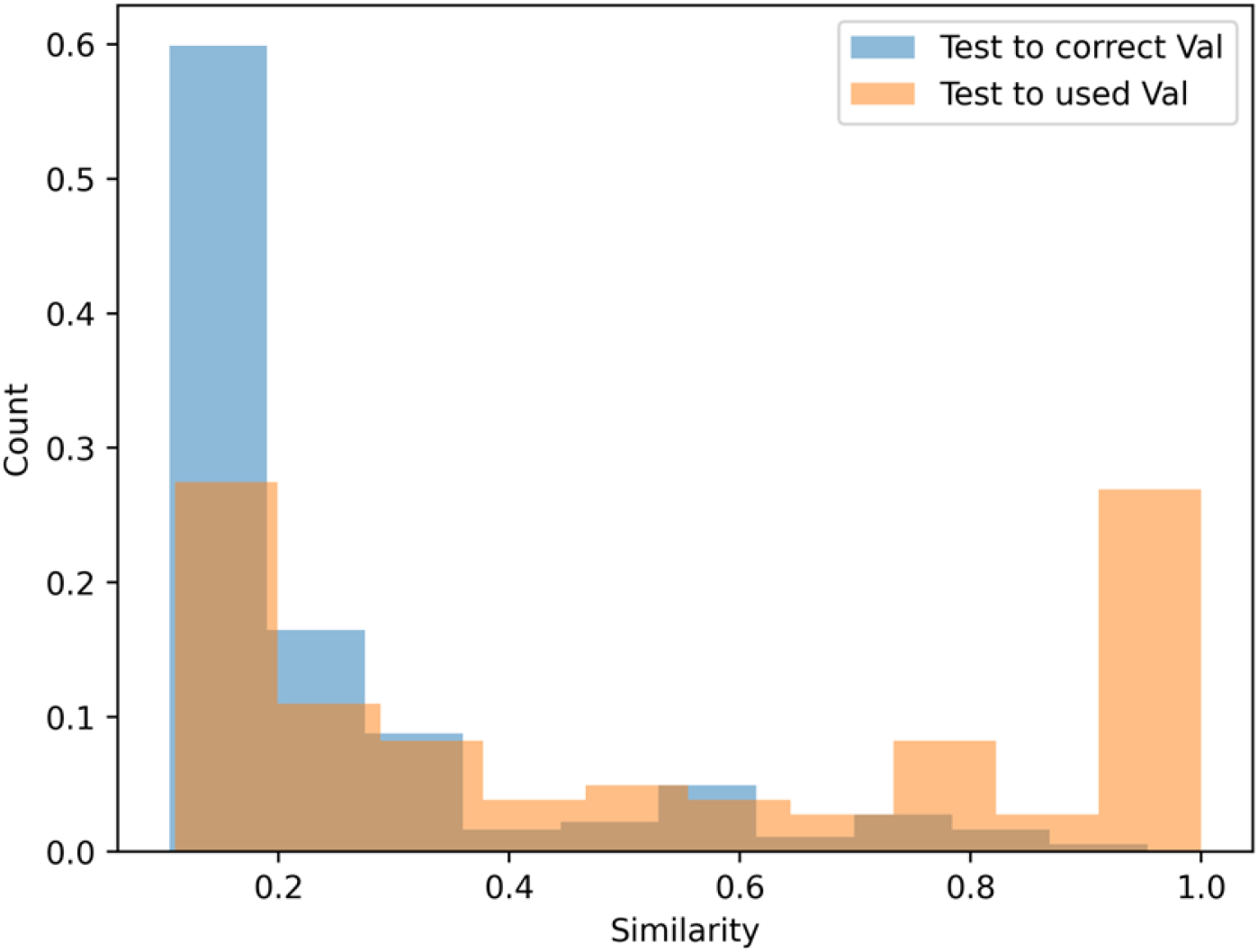
Distribution of molecules, which have different similarities, between TDC *caco2* test set and both the validation set used during tuning and the correctly generated validation set.

Although the core model training procedures are implemented correctly in both cases, the data leakage at the hyperparameters optimization leads to inflated reported leaderboard metrics. After correcting the split function and re-evaluating the models we obtained substantially lower leaderboard ranks— third place for GradientBoost (decrease from first) and eighth place for XGBoost (decrease from second).

### Re-evaluation of model performance

Only three models reached the final testing stage: **CaliciBoost** (first on the *caco2_wang*), **MapLight + GNN** (first on *clearance_microsome_az, cyp2c9_veith, cyp2d6_veith, cyp3a4_veith, herg, pgp_broccatelli*, and *vdss_lombardo*), and **MapLight**.

CaliciBoost was developed exclusively for the evaluation of Caco-2 effective permeability. Our MAE estimate for CaliciBoost was 0.271 ± 0.002, compared with 0.256 ± 0.006 reported in TDC, while the model retained its first-place ranking. The small discrepancy between these values is most likely related to the updates of the TDC datasets, which are, unfortunately, not under version control. That is why it is not possible to attribute the benchmarking results to a particular well-defined dataset version unambiguously. Part of observed discrepancies may also be related to differences in computational hardware and/or software environments used for evaluation.

MapLight and MapLight + GNN were evaluated across all 22 TDC ADMET endpoints. A summary of their re-evaluated performance is shown in Table 1.

**Table 2.** Original and re-evaluated performance metrics and ranks of Maplight and Maplight+ GNN models on different TDC ADMET endpoints.

| Task | Maplight |  |  |  | Maplight+ GNN |  |  |  |
| --- | --- | --- | --- | --- | --- | --- | --- | --- |
|  | TDC metric | TDC rank | reproduced metric | reproduced rank | TDC metric | TDC rank | reproduced metric | reproduced rank |
| Caco2 | $0.276 \pm 0.005$ | 3 | $0.276 \pm 0.006$ | 3 | $0.287 \pm 0.005$ | 6 | $0.29 \pm 0.008$ | 7 |
| Bio | $0.930 \pm 0.002$ | 4 | $0.741 \pm 0.007$ | -- | $0.938 \pm 0.002$ | 2 | $0.74 \pm 0.01$ | -- |
| Lipo | $0.539 \pm 0.002$ | 10 | $0.542 \pm 0.003$ | 11 | $0.525 \pm 0.003$ | 7 | $0.525 \pm 0.003$ | 7 |
| Solu | $0.792 \pm 0.002$ | 6 | $0.787 \pm 0.006$ | 5 | $0.789 \pm 0.003$ | 5 | $0.795 \pm 0.004$ | 6 |
| HIA | $0.986 \pm 0.000$ | 5 | $0.983 \pm 0.001$ | 6 | $0.989 \pm 0.001$ | 3 | $0.989 \pm 0.001$ | 3 |
| Pgp | $0.930 \pm 0.002$ | 3 | $0.932 \pm 0.001$ | 3 | $0.938 \pm 0.002$ | 1 | $0.938 \pm 0.0$ | 1 |
| BBB | $0.916 \pm 0.001$ | 3 | $0.92 \pm 0.002$ | 2 | $0.913 \pm 0.001$ | 5 | $0.916 \pm 0.002$ | 4 |
| PPBR | $7.660 \pm 0.058$ | 3 | $7.622 \pm 0.065$ | 3 | $7.526 \pm 0.106$ | 2 | $7.573 \pm 0.163$ | 2 |
| VDss | $0.707 \pm 0.009$ | 2 | $0.716 \pm 0.005$ | 1 | $0.713 \pm 0.007$ | 1 | $0.716 \pm 0.006$ | 2 |
| CYP2c9 Veith | $0.783 \pm 0.002$ | 7 | $0.786 \pm 0.004$ | 7 | $0.859 \pm 0.001$ | 1 | $0.859 \pm 0.001$ | 1 |
| CYP2d6 Veith | $0.723 \pm 0.003$ | 3 | $0.72 \pm 0.002$ | 5 | $0.790 \pm 0.001$ | 1 | $0.79 \pm 0.001$ | 1 |
| CYP3a4 Veith | $0.881 \pm 0.001$ | 5 | $0.881 \pm 0.001$ | 5 | $0.916 \pm 0.000$ | 1 | $0.916 \pm 0.0$ | 1 |
| CYP2c9 Sub | $0.415 \pm 0.008$ | 8 | $0.411 \pm 0.005$ | 8 | $0.437 \pm 0.008$ | 3 | $0.446 \pm 0.015$ | 2 |
| **CYP2d6 Sub** | 0.713 ± 0.009 | 4 | 0.718 ± 0.008 | 4 | 0.720 ± 0.002 | 3 | 0.716 ± 0.007 | 3 |
| **CYP3a4 Sub** | 0.650 ± 0.006 | 5 | 0.649 ± 0.006 | 5 | 0.647 ± 0.008 | 6 | 0.642 ± 0.004 | 6 |
| **Half** | 0.562 ± 0.008 | 2 | 0.559 ± 0.011 | 3 | 0.557 ± 0.034 | 3 | 0.563 ± 0.009 | 2 |
| **Clearenc e_hepa** | 0.466 ± 0.012 | 3 | 0.469 ± 0.016 | 3 | 0.498 ± 0.009 | 2 | 0.5 ± 0.012 | 2 |
| **Clearenc e_micro** | 0.626 ± 0.008 | 3 | 0.625 ± 0.009 | 5 | 0.630 ± 0.010 | 1 | 0.635 ± 0.007 | 1 |
| **LD50_Zh u** | 0.621 ± 0.003 | 7 | 0.618 ± 0.005 | 7 | 0.633 ± 0.003 | 12 | 0.629 ± 0.005 | 10 |
| **hERG** | 0.871 ± 0.004 | 4 | 0.873 ± 0.004 | 4 | 0.880 ± 0.002 | 1 | 0.88 ± 0.003 | 1 |
| **AMES** | 0.868 ± 0.002 | 4 | 0.865 ± 0.001 | 4 | 0.869 ± 0.002 | 3 | 0.872 ± 0.001 | 1 |
| **DILI** | 0.887 ± 0.006 | 10 | 0.883 ± 0.003 | 12 | 0.917 ± 0.005 | 5 | 0.918 ± 0.005 | 5 |

reproduced rank got better. reproduced rank got worse. reproduced rank got much worse

It is evident that MapLight+GNN retains its leaderboard rank in 13 out of 22 ADMET endpoints, improves its position in five cases (notably reaching first place for *ames*), and exhibits a lower rank in three cases. A substantial decline in performance is observed only for *bioavailability_ma*.

Similarly, MapLight preserves its original rank in 11 endpoints, improves its ranking in four cases (including attaining first place for *vdss_lombardo*), and shows a lower rank in six cases. As with MapLight+GNN, a pronounced drop in performance is again observed for *bioavailability_ma*.

Across all 22 endpoints, the absolute differences between TDC-reported and reproduced metrics are very small (< 0.01) for the vast majority of cases for both models, indicating high overall reproducibility of the leaderboard results. For most ADMET tasks, reproduced metrics fall within overlapping uncertainty ranges of the TDC values, and rank changes (where present) are small and consistent with the narrow performance gaps among top-ranked models. Minor deviations are likely attributable to routine technical factors such as random seed variability, numerical precision, or differences in software and hardware environments used for evaluation. It should also be noted that the TDC platform does not provide explicit dataset or leaderboard versioning. As a result, minor changes in dataset composition, preprocessing, or split definitions over time may contribute to small discrepancies between reproduced and leaderboard metrics, even when identical evaluation protocols are followed.

In general, MapLight+GNN tends to outperform MapLight, particularly for metabolism-related endpoints (*cyp2c9_veith, cyp2d6_veith, cyp3a4_veith*, and *clearance_microsome_az*), where MapLight+GNN consistently occupies top leaderboard positions and maintains close agreement between reproduced and TDC metrics. For several endpoints — including *hERG, pgp_broccatelli, and cyp3a4_veith* — the reproduced ranks for both models closely match the TDC leaderboard positions, highlighting stable and reproducible performance in these tasks.

The largest and systematic discrepancies between TDC and reproduced results are consistently observed for *bioavailability_ma*, where both models exhibit markedly lower reproduced AUROC values and substantial rank changes. A second, but less pronounced, deviation is observed for *ppbr_az*, where MAE differences are larger than for other endpoints, although rankings remain comparatively stable. Given that TDC doesn’t maintain publicly versioned dataset releases, it is possible that the benchmark snapshot used to obtain the original leaderboard results differs from the dataset currently available. Such untracked updates (e.g., removal/addition of molecules or adjustments to train-test splits) could contribute to systematic shifts in performance for particular endpoints.

### Performance testing of in-house models

The summarized results of our in-house model performance evaluation across 22 ADMET endpoints are presented in Table S3. For the majority of endpoints, the two-stage optimization procedure — Sequential Forward Selection (SFS) followed by Bayesian hyperparameter optimization (HPO) — leads to improved predictive performance compared with the baseline models, as reflected in better evaluation metrics and/or higher leaderboard rankings. This indicates that, in most cases, systematic feature selection combined with targeted hyperparameter tuning is an effective strategy for enhancing model performance within the LightGBM framework. Notable exceptions are:

(a) *dili a*nd *pgp_broccatelli*, for which the baseline variants of the models remain the most competitive;
(b) *bbb_martins, cyp3a4_substrate_carbonmangels, clearance_hepatocyte_az, clearance_microsome_az*, and *ames*, for which the models after the first optimization stage (SFS) achieve the best performance among the three tested variants. In the case of *bbb_martins*, SFS substantially improves the model position from 8th to 2nd place on the leaderboard, whereas subsequent HPO results in a marked drop in ranking despite comparable or slightly improved metrics.

For *ppbr_az* and *half_life_obach*, performance temporarily deteriorates after SFS but improves after HPO, ultimately yielding better metrics and rankings than those of the corresponding baseline models. This suggests that, for these endpoints, feature selection alone is insufficient, whereas hyperparameter tuning is critical for achieving optimal performance.

In the case of *hia_hou*, the optimized model reaches first place on the corresponding leaderboard. In addition, the optimized models for *ld50_zhu, lipophilicity_astrazeneca, solubility_aqsoldb*, and *cyp2c9_veith* rank within the top five positions of their respective leaderboards, indicating consistently strong performance across both physicochemical and metabolism-related tasks.

A notable pattern emerges for several binary classification endpoints with very high reported AUROC values on the TDC leaderboards. In *hia_hou, bbb_martins*, and *pgp_broccatelli*, a substantial fraction of leaderboard models exhibit AUROC values above 0.9 (17 out of 25 models for *hia_hou*, 12 out of 25 for *bbb_martins*, and 9 out of 25 for *pgp_broccatelli*). The AUROC values obtained for our optimized models on these endpoints also fall within this high-performance range. This suggests that achieving AUROC values above 0.9 on these benchmarks should not automatically be interpreted as evidence of overfitting and can be regarded as a normal performance regime for these tasks. However, this does not rule out the presence of overfitted models in the TDC ADMET leaderboards.

### The effect of deliberate model overfitting

In order to study the effect of possible unintentional or deliberate model overfitting to the test set we constructed deliberately overfitted versions of our in-house models and analyzed their ranking patterns in comparison with those of other top-performing models such as MapLight+GNN and MiniMol.

Table S4 summarizes the performance of deliberately overfitted in-house models across all 22 TDC ADMET endpoints. In contrast to the “honest” models, the overfitted variants were intentionally tuned to maximize performance on the public TDC test sets, allowing us to examine how sensitive leaderboard rankings are to extreme, test-set–driven optimization. For each endpoint, we consider the best achieved leaderboard position across the three variants (baseline, SFS, and HPO).

A clear and systematic pattern emerges. For a substantial fraction of benchmarks, deliberate overfitting leads to marked improvements in leaderboard positions, often elevating models from the middle or lower tiers directly into the top ranks. The most pronounced effects are observed for *bbb_martins* (rank 8 → 1), *clearance_microsome_az* (14 → 1), *cyp2d6_substrate* (12 → 1), and *ames* (13 → 1). Similarly strong gains are seen for *caco2_wang* (9 → 2), *cyp3a4_substrate* (14 → 2), *dili* (13 → 2), and *hia_hou* (2 → 1).

For another group of endpoints, overfitting improves performance more moderately, consistently moving models upward in the leaderboard without necessarily reaching first place. This behavior is observed for *lipophilicity_astrazeneca* (13 → 5), *solubility_aqsoldb* (7 → 3), *ld50_zhu* (6 → 3), *half_life_obach* (11 → 9), *and ppbr_az* (9 → 5, then 7 after HPO).

In contrast, bioavailability_ma remains largely insensitive to deliberate overfitting: all three variants (baseline, SFS, and HPO) retain rank 21 despite noticeable changes in AUROC values. This likely indicates that our model architecture is generally unsuitable for this endpoint. On the other hand, the issue may originate from the benchmark itself. Specifically, it is possible that the dataset was updated without public disclosure, resulting in the inability to reproduce the leaderboard metrics reported for the top-ranked models before (see our verification results for MapLight + GNN and MapLight, both of which failed to reproduce the reported performance on this endpoint).

Quantitatively, deliberate overfitting produces a substantial upward shift in leaderboard positions: the overfitted model reaches 1st place in 5/22 endpoints, 2nd place in 3/22, and 3rd place in 2/22, i.e., it appears in the top three in 10/22 endpoints overall (≈45%) (table 2). In contrast, our conventionally trained (“honest”) in-house models reach the top three only twice across the same 22 endpoints (best rank across baseline/SFS/HPO), highlighting a pronounced gap between standard optimization and test-set–driven tuning.

**Table 2.** Best metrics and ranks of in-house models across different TDC ADMET endpoints

| **benchmark** | **metric name** | **honest metric** | **honest rank** | **overfitted metric** | **overfitted rank** |
| --- | --- | --- | --- | --- | --- |
| caco2_wang | MAE | 0.2875 | 7 | 0.2694 | 2 |
| bioavailability_ma | AUROC | 0.6744 | 21 | 0.7147 | 21 |
| lipophilicity_astrazeneca | MAE | 0.5108 | 5 | 0.5112 | 5 |
| solubility_aqsolddb | MAE | 0.7808 | 5 | 0.7735 | 3 |
| hia_hou | AUROC | 0.9938 | 1 | 0.9951 | 1 |
| pgp_broccatelli | AUROC | 0.902 | 10 | 0.9202 | 7 |
| bbb_martins | AUROC | 0.9225 | 2 | 0.9376 | 1 |
| ppbr_az | MAE | 7.9457 | 7 | 7.779 | 7 |
| vdss_lombardo | Spearman | 0.4949 | 11 | 0.4909 | 13 |
| cyp2c9_veith | AUPRC | 0.7922 | 5 | 0.792 | 5 |
| cyp2d6_veith | AUPRC | 0.7178 | 7 | 0.7085 | 5 |
| cyp3a4_veith | AUPRC | 0.8748 | 9 | 0.8826 | 5 |
| cyp2c9_substrate_carbonmangels | AUPRC | 0.3459 | 19 | 0.3566 | 18 |
| cyp2d6_substrate_carbonmangels | AUPRC | 0.6842 | 10 | 0.7566 | 1 |
| cyp3a4_substrate_carbonmangels | AUROC | 0.6384 | 9 | 0.6658 | 2 |
| half_life_obach | Spearman | 0.3927 | 9 | 0.4089 | 9 |
| clearance_hepatocyte_az | Spearman | 0.3353 | 16 | 0.4413 | 6 |
| clearance_microsome_az | Spearman | 0.5542 | 14 | 0.6314 | 1 |
| ld50_zhu | MAE | 0.5866 | 4 | 0.5766 | 3 |
| herg | AUROC | 0.8032 | 11 | 0.8495 | 6 |
| ames | AUROC | 0.8422 | 10 | 0.8723 | 1 |
| dili | AUROC | 0.86 | 16 | 0.9422 | 2 |

Considering the TDC ADMET leaderboards at the time of the study, the top-three ranks are dominated by a small set of recurring models rather than a large, diverse pool of methods. Among these, MapLight appears in the top three 17 times (1 first place, 5 seconds, and 11 thirds), MapLight+GNN 15 times (7 firsts, 5 seconds, and 3 thirds), and MiniMol 13 times (7 firsts, 4 seconds, and 2 thirds), with additional recurrent contributions from CFA (8 appearances), ZairaChem (5), and ADMETrix (4).

Our deliberately overfitted model reaches the top three in 10 out of 22 endpoints, demonstrating that test-set–driven tuning can elevate leaderboard positions but can not dominate all leaderboards. By comparison, MapLight (17/22), MapLight+GNN (15/22), and MiniMol (13/22) appear in the top three substantially more often than any of our in-house overfitted models tested here. This suggests that the recurrent TDC leaders are likely to be superior in the choice of model architecture, descriptor set or training pipeline, rather than by deliberate fitting to a specific endpoint.

## Discussion

The present study reveals systematic limitations in using the open TDC leaderboard as a direct indicator of ADMET model quality. Our re-evaluation of top-ranked TDC models shows that full reproducibility is achievable for only three methods - CaliciBoost, MapLight and MapLight+GNN, while all other top-ranked models are prone to serious technical or methodological issues. At the same time, our experiments with deliberately overfitted in-house models demonstrate that accidental or deliberate model tuning to the public test set can substantially alter leaderboard positions, often elevating otherwise mediocre models to top ranks.

Some models that occupied the TDC ADMET top-3 at the time of writing could not be reliably used in practice due to unavailable source code, impossibility to set up an execution environment, inability to run the model in a properly configured environment or critical flaws in data processing and optimization procedures. This points to a structural weakness in the current leaderboard format: high ranks are not accompanied by systematic verification of code availability, functionality, and methodological correctness.

An analysis of the model repositories showed that reproducibility issues in many cases stem from mistakes and omissions in the installation and running instructions. This signifies insufficient scrutiny in software development and distribution practices, which is, unfortunately, common in academic software development in general and in the ML field in particular. The availability of a public repository alone does not guarantee the practical reproducibility of models in the absence of complete, well-defined and well-tested installation instructions.

Our analysis shows that although only about 30% of the selected models reached the final stage of verification, this subset overall demonstrates good reproducibility. For MapLight, MapLight+GNN, and CaliciBoost, discrepancies between reported and reproduced metrics were small and plausibly attributable to technical factors (differences in hardware, software environments, or library versions). This confirms that, when implemented correctly, top TDC models can be reproducible, but simultaneously highlights that a substantial fraction of leaders does not meet this standard.

Comparison of our in house “honest” models with the ranges of TDC metrics shows that their values consistently fall within the characteristic ranges of the corresponding benchmarks. This indicates that the metric ranges reported on the leaderboards are broadly realistic and are not merely a consequence of pervasive overfitting.

However, a markedly different picture emerges when analyzing the sensitivity of leaderboards to overfitting. Our deliberately overfitted models entered the top-3 in 10 out of 22 endpoints, whereas “honest” models did so in only 2 out of 22. This contrast demonstrates that public leaderboards are highly susceptible to accidental or deliberate overfitting to the open testing dataset.

At the same time, leading TDC models (MapLight - 17/22, MapLight+GNN - 15/22, MiniMol - 13/22 top-3 appearances) appear in the top ranks far more frequently than any of our own overfitted variants. Such dominance in a broad range of endpoints is difficult to reproduce within a single model architecture even under deliberate overfitting. Such dominance across a wide range of endpoints cannot be easily explained by tuning for specific endpoints, as models such as MapLight use a fixed set of descriptors that is the same for all tasks. Rather, this consistency suggests that the selected descriptors represent molecular properties that influence multiple ADMET processes, allowing the same modeling pipeline to perform well across different endpoints. The model authors do not disclose how exactly this particular set of descriptors was selected, but in terms of model performance, it appears to be quite successful.

These conclusions align with reports indicating that long-lived open test sets encourage gradual adaptation of models to a specific benchmark, even without deliberate “cheating” (45,46). Moreover, in some cases the reported accuracy may exceed the reproducibility of the underlying experimental measurements themselves, which can serve as an indicator of hidden overfitting (30,31).

One of the crucial findings of this study is the absence of exact reproducibility of the TDC benchmarks even for the perfectly described models. We believe that it originates from the absence of explicit dataset versioning within the TDC benchmark suite. The platform does not provide fixed dataset releases, version identifiers, check sums or detailed changelogs describing updates to data composition and/or processing, so the dataset snapshot used to generate leaderboard metrics may differ from the version available at the time of reproduction. Consequently, some discrepancies between reproduced and leaderboard results may reflect dataset evolution rather than genuine methodological advantages or limitations of the models. More broadly, the lack of transparent versioning also reduces the practical usability of the benchmark, as it complicates cross-study comparisons and makes it difficult to determine whether performance discrepancies arise from the modeling pipeline features or from undocumented changes in the underlying data.

TDC occupies a distinctive leading position among publicly available resources for ADMET prediction benchmarking, while other platforms implement somewhat different benchmarking models. DREAM (47) and SAMPL (48) rely primarily on time-limited blind challenges with centralized evaluation on hidden data; CodaBench (49) offers infrastructure for containerized benchmarks but does not maintain its own ADMET leaderboard; MoleculeNet (50) standardizes datasets and metrics but does not run a continuous ranking; PharmaBench (51) emphasizes reproducible workflows but lacks a community-wide public leaderboard; OpenADMET (52) maintains an open repository of ADMET models while developing supporting tools for training and model-to-model comparison. Therefore, none of these alternatives combines broad ADMET coverage with a continuously maintained public leaderboard in the way TDC does.

At the same time TDC leaderboards also demonstrate significant shortcomings, which have to be addressed in the next generation of ADMET benchmarks:

- The test set should not be publicly available to avoid deliberate or accidental model fitting towards it.
- The dataset should be strictly versioned with a distinct checksum assigned to each release or snapshot. Reported model results should be tied to this checksum to allow exact reproducibility.
- The model authors should submit the models rather than their results, complete with the inference environments in a standardised format. This will allow automatic quality control, reproducibility and proper versioning of the leaderboard.

Taken together, our results indicate that TDC leaderboards are useful as a reference point, but high positions should not be interpreted as direct evidence of a model’s predictive strength.

Under conditions of fully open test data, TDC leaderboard results can be highly sensitive to various forms of data leakage, including apparent or hidden optimization of models for a specific benchmark sample.

## Author contributions

AN served as the conceptual lead and originator of the core idea. IK, RS, NS and MM performed TDC models assessment, developed in-house models and performed the analysis. SY, SS and VH coordinated the work, critically assessed the results, and guided the analysis. SS, SY and AN designed the study and controlled its progress. The manuscript was written by ON and SY.

## Acknowledgements

SY has received funding through the grant MSMT-355/2025-16 from the Ministry of Education, Youth and Sport of the Czech Republic.

The authors acknowledge the use of the large language model GPT-5.2 for proofreading and stylistic editing of the manuscript.

## References

1. DiMasi JA, Grabowski HG, Hansen RW. Innovation in the pharmaceutical industry: New estimates of R&D costs. J Health Econ. 2016 May 1;47:20–33. doi:10.1016/j.jhealeco.2016.01.012

2. Wouters OJ, McKee M, Luyten J. Estimated Research and Development Investment Needed to Bring a New Medicine to Market, 2009-2018. JAMA. 2020 Mar 3;323(9):844–53. doi:10.1001/jama.2020.1166

3. Cheng F, Li W, Liu G, Tang Y. In Silico ADMET Prediction: Recent Advances, Current Challenges and Future Trends. http://www.eurekaselect.com [Internet]. [cited 2026 Feb 20]. Available from: https://www.eurekaselect.com/article/52804

4. Kola I, Landis J. Can the pharmaceutical industry reduce attrition rates? Nat Rev Drug Discov. 2004 Aug;3(8):711–6. doi:10.1038/nrd1470

5. Waring MJ, Arrowsmith J, Leach AR, Leeson PD, Mandrell S, Owen RM, et al. An analysis of the attrition of drug candidates from four major pharmaceutical companies. Nat Rev Drug Discov. 2015 Jul;14(7):475–86. doi:10.1038/nrd4609

6. Ferreira LLG, Andricopulo AD. ADMET modeling approaches in drug discovery. Drug Discov Today. 2019 May 1;24(5):1157–65. doi:10.1016/j.drudis.2019.03.015

7. Daina A, Michielin O, Zoete V. SwissADME: a free web tool to evaluate pharmacokinetics, drug-likeness and medicinal chemistry friendliness of small molecules. Sci Rep. 2017 Mar 3;7(1):42717. doi:10.1038/srep42717

8. Huang K, Fu T, Glass LM, Zitnik M, Xiao C, Sun J. DeepPurpose: a deep learning library for drug–target interaction prediction. Bioinformatics. 2020 Dec 12;36(22–23):5545–7. doi:10.1093/bioinformatics/btaa1005 PubMed PMID: 33275143; PubMed Central PMCID: PMC8016467.

9. Lipinski CA, Lombardo F, Dominy BW, Feeney PJ. Experimental and computational approaches to estimate solubility and permeability in drug discovery and development settings. Adv Drug Deliv Rev. 1997 Jan 15;In Vitro Models for Selection of Development Candidates 23(1):3–25. doi:10.1016/S0169-409X(96)00423-1

10. Vamathevan J, Clark D, Czodrowski P, Dunham I, Ferran E, Lee G, et al. Applications of machine learning in drug discovery and development. Nat Rev Drug Discov. 2019 Jun;18(6):463–77. doi:10.1038/s41573-019-0024-5

11. He J, Nittinger E, Tyrchan C, Czechtizky W, Patronov A, Bjerrum EJ, et al. Transformer-based molecular optimization beyond matched molecular pairs. J Cheminformatics. 2022 Mar 28;14(1):18. doi:10.1186/s13321-022-00599-3

12. Fan N, Chen J, Wang J, Chen ZS, Yang Y. Bridging data and drug development: Machine learning approaches for next-generation ADMET prediction. Drug Discov Today. 2025 Nov 1;30(11):104487. doi:10.1016/j.drudis.2025.104487

13. Pathan I, Raza A, Sahu A, Joshi M, Sahu Y, Patil Y, et al. Revolutionizing pharmacology: AI-powered approaches in molecular modeling and ADMET prediction. Med Drug Discov. 2025 Dec 1;28:100223. doi:10.1016/j.medidd.2025.100223

14. Gaudelet T, Day B, Jamasb AR, Soman J, Regep C, Liu G, et al. Utilizing graph machine learning within drug discovery and development. Brief Bioinform. 2021 Nov 1;22(6):bbab159. doi:10.1093/bib/bbab159

15. Wang Y, Wang J, Cao Z, Barati Farimani A. Molecular contrastive learning of representations via graph neural networks. Nat Mach Intell. 2022 Mar;4(3):279–87. doi:10.1038/s42256-022-00447-x

16. Vishwakarma G, Sonpal A, Hachmann J. Metrics for Benchmarking and Uncertainty Quantification: Quality, Applicability, and Best Practices for Machine Learning in Chemistry. Trends Chem. 2021 Feb 1;Special Issue: Machine Learning for Molecules and Materials 3(2):146–56. doi:10.1016/j.trechm.2020.12.004

17. Bender A, Schneider N, Segler M, Patrick Walters W, Engkvist O, Rodrigues T. Evaluation guidelines for machine learning tools in the chemical sciences. Nat Rev Chem. 2022 Jun;6(6):428–42. doi:10.1038/s41570-022-00391-9

18. Artrith N, Butler KT, Coudert FX, Han S, Isayev O, Jain A, et al. Best practices in machine learning for chemistry. Nat Chem. 2021 Jun;13(6):505–8. doi:10.1038/s41557-021-00716-z

19. Yang K, Swanson K, Jin W, Coley C, Eiden P, Gao H, et al. Analyzing Learned Molecular Representations for Property Prediction. J Chem Inf Model. 2019 Aug 26;59(8):3370–88. doi:10.1021/acs.jcim.9b00237

20. Kumar N, Acharya V. Machine intelligence-driven framework for optimized hit selection in virtual screening. J Cheminformatics. 2022 Jul 22;14(1):48. doi:10.1186/s13321-022-00630-7

21. Zhao ZW, Cueto M del, Troisi A. Limitations of machine learning models when predicting compounds with completely new chemistries: possible improvements applied to the discovery of new non-fullerene acceptors [Internet]. 2022 Jun 13. doi:10.1039/D2DD00004K

22. Wu Z, Ramsundar B, Feinberg EN, Gomes J, Geniesse C, Pappu AS, et al. MoleculeNet: a benchmark for molecular machine learning. Chem Sci. 2018 Jan 3;9(2):513–30. doi:10.1039/C7SC02664A

23. Niu Z, Xiao X, Wu W, Cai Q, Jiang Y, Jin W, et al. PharmaBench: Enhancing ADMET benchmarks with large language models. Sci Data. 2024 Sep 10;11(1):985. doi:10.1038/s41597-024-03793-0

24. Huang K, Fu T, Gao W, Zhao Y, Roohani Y, Leskovec J, et al. Therapeutics Data Commons: Machine Learning Datasets and Tasks for Drug Discovery and Development [Internet]. arXiv; 2021 [cited 2026 Feb 20]. Available from: http://arxiv.org/abs/2102.09548 doi:10.48550/arXiv.2102.09548

25. Huang K, Fu T, Gao W, Zhao Y, Roohani Y, Leskovec J, et al. Artificial intelligence foundation for therapeutic science. Nat Chem Biol. 2022 Oct;18(10):1033–6. doi:10.1038/s41589-022-01131-2 PubMed PMID: 36131149; PubMed Central PMCID: PMC9529840.

26. Velez-Arce A, Huang K, Li MM, Lin X, Gao W, Fu T, et al. TDC-2: Multimodal Foundation for Therapeutic Science [Internet]. bioRxiv; 2024 [cited 2026 Feb 20]. p. 2024.06.12.598655. Available from: https://www.biorxiv.org/content/10.1101/2024.06.12.598655v1 doi:10.1101/2024.06.12.598655

27. Kapoor S, Narayanan A. Leakage and the reproducibility crisis in machine-learning-based science. Patterns. 2023 Sep 8;4(9). doi:10.1016/j.patter.2023.100804 PubMed PMID: 37720327.

28. Apicella A, Isgrò F, Prevete R. Don’t push the button! Exploring data leakage risks in machine learning and transfer learning. Artif Intell Rev. 2025 Aug 20;58(11):339. doi:10.1007/s10462-025-11326-3

29. Mania H, Miller J, Schmidt L, Hardt M, Recht B. Model Similarity Mitigates Test Set Overuse [Internet]. arXiv; 2019 [cited 2026 Feb 20]. Available from: http://arxiv.org/abs/1905.12580 doi:10.48550/arXiv.1905.12580

30. Crusius D, Cipcigan F, C. Biggin P. Are we fitting data or noise? Analysing the predictive power of commonly used datasets in drug-, materials-, and molecular-discovery [Internet]. 2025 Jan 16. doi:10.1039/D4FD00091A

31. Saidi P, Dasarathy G, Berisha V. Unraveling overoptimism and publication bias in ML-driven science. Patterns. 2025 Apr 11;6(4):101185. doi:10.1016/j.patter.2025.101185

32. Ash JR, Wognum C, Rodríguez-Pérez R, Aldeghi M, Cheng AC, Clevert DA, et al. Practically Significant Method Comparison Protocols for Machine Learning in Small Molecule Drug Discovery. J Chem Inf Model. 2025 Sep 22;65(18):9398–411. doi:10.1021/acs.jcim.5c01609

33. Gadaleta D, Serrano-Candelas E, Ortega-Vallbona R, Colombo E, Garcia de Lomana M, Biava G, et al. Comprehensive benchmarking of computational tools for predicting toxicokinetic and physicochemical properties of chemicals. J Cheminformatics. 2024 Dec 26;16(1):145. doi:10.1186/s13321-024-00931-z

34. Bender A, Cortés-Ciriano I. Artificial intelligence in drug discovery: what is realistic, what are illusions? Part 1: Ways to make an impact, and why we are not there yet. Drug Discov Today. 2021 Feb 1;26(2):511–24. doi:10.1016/j.drudis.2020.12.009

35. Bajorath J, Kearnes S, Walters WP, Meanwell NA, Georg GI, Wang S. Artificial Intelligence in Drug Discovery: Into the Great Wide Open. J Med Chem. 2020 Aug 27;63(16):8651–2. doi:10.1021/acs.jmedchem.0c01077

36. Scalia G, Grambow CA, Pernici B, Li YP, Green WH. Evaluating Scalable Uncertainty Estimation Methods for Deep Learning-Based Molecular Property Prediction. J Chem Inf Model. 2020 Jun 22;60(6):2697–717. doi:10.1021/acs.jcim.9b00975

37. lightgbm.LGBMRegressor — LightGBM 4.6.0.99 documentation [Internet]. [cited 2026 Feb 20]. Available from: https://lightgbm.readthedocs.io/en/latest/pythonapi/lightgbm.LGBMRegressor.html

38. lightgbm.LGBMClassifier — LightGBM 4.6.0.99 documentation [Internet]. [cited 2026 Feb 20]. Available from: https://lightgbm.readthedocs.io/en/latest/pythonapi/lightgbm.LGBMClassifier.html

39. Akiba T, Sano S, Yanase T, Ohta T, Koyama M. Optuna: A Next-generation Hyperparameter Optimization Framework. In: Proceedings of the 25th ACM SIGKDD International Conference on Knowledge Discovery & Data Mining [Internet]. New York, NY, USA: Association for Computing Machinery; 2019 [cited 2026 Feb 20]. p. 2623–31. (KDD ‘19). Available from: https://dl.acm.org/doi/10.1145/3292500.3330701 doi:10.1145/3292500.3330701

40. Watanabe S. Tree-Structured Parzen Estimator: Understanding Its Algorithm Components and Their Roles for Better Empirical Performance [Internet]. arXiv; 2025 [cited 2026 Feb 20]. Available from: http://arxiv.org/abs/2304.11127 doi:10.48550/arXiv.2304.11127

41. jiang N, Quazi M, Schweikert C, Hsu DF, Oprea T, sirimulla suman. Enhancing ADMET Property Models Performance through Combinatorial Fusion Analysis. ChemRxiv. 2023(1129). doi:10.26434/chemrxiv-2023-dh70x

42. partex-nv-opensource. GitHub [Internet]. [cited 2026 Feb 20]. tdc-submission/src/ics-admet-prediction-tdc-versiontwo/report/Advancing ADMET Prediction_A Hybrid Machine Learning Framework (Partex ADMETrix).pdf at main · partex-nv-opensource/tdc-submission. Available from: https://github.com/partex-nv-opensource/tdc-submission/blob/main/src/ics-admet-prediction-tdc-versiontwo/report/Advancing%20ADMET%20Prediction_%20A%20Hybrid%20Machine%20Learning%20Framework%20(Partex%20ADMETrix).pdf

43. KatanaGraph. GitHub [Internet]. [cited 2026 Feb 20]. SimGCN-TDC/Report_SimGCN_for_TDC_Benchmarks.pdf at main · KatanaGraph/SimGCN-TDC. Available from: https://github.com/KatanaGraph/SimGCN-TDC/blob/main/Report_SimGCN_for_TDC_Benchmarks.pdf

44. Turon G, Hlozek J, Woodland JG, Chibale K, Duran-Frigola M. First fully-automated AI/ML virtual screening cascade implemented at a drug discovery centre in Africa [Internet]. bioRxiv; 2022 [cited 2026 Feb 20]. p. 2022.12.13.520154. Available from: https://www.biorxiv.org/content/10.1101/2022.12.13.520154v1 doi:10.1101/2022.12.13.520154

45. Blum A, Hardt M. The Ladder: A Reliable Leaderboard for Machine Learning Competitions. In: Proceedings of the 32nd International Conference on Machine Learning [Internet]. PMLR; 2015 [cited 2026 Feb 20]. p. 1006–14. Available from: https://proceedings.mlr.press/v37/blum15.html

46. Kapoor S, Narayanan A. Leakage and the Reproducibility Crisis in ML-based Science [Internet]. arXiv; 2022 [cited 2026 Feb 20]. Available from: http://arxiv.org/abs/2207.07048 doi:10.48550/arXiv.2207.07048

47. Aaron. DREAM Challenges [Internet]. [cited 2026 Feb 20]. DREAM. Available from: https://dreamchallenges.org/

48. Mobley DL. SAMPL [Internet]. [cited 2026 Feb 20]. SAMPL challenges. Available from: https://samplchallenges.org//

49. Xu Z, Escalera S, Pavão A, Richard M, Tu WW, Yao Q, et al. Codabench: Flexible, easy-to-use, and reproducible meta-benchmark platform. Patterns. 2022 Jul;3(7):100543. doi:10.1016/j.patter.2022.100543

50. Pande Group. MoleculeNet [Internet]. [cited 2026 Feb 20]. Available from: https://moleculenet.org/

51. mindrank-ai/PharmaBench [Jupyter Notebook] [Internet]. MindRank Technologies; 2026 [cited 2026 Feb 20]. Available from: https://github.com/mindrank-ai/PharmaBench

52. OMSF [Internet]. [cited 2026 Feb 20]. OpenADMET. Available from: https://openadmet.org/

