## Supplementary Information for "Critical Assessment of ML models for ADMET Prediction in TDC leaderboards"

### Supplementary Materials

*Ihor Koleiev [1,2], Roman Stratiichuk [1,3], Nazar Shevchuk [1], Mykola Melnychenko [1], Oleksiy Nyporko, Daniil Todoryshyn [1], Vladyslav Husak [1,4], Sergii Starosyla [1], Semen Yesylevskyy [1,2,5,6,*], Alan Nafiiev [1].*

[1] Receptor.AI Inc., 20-22 Wenlock Road, London N1 7GU, United Kingdom.

[2] Department of Physics of Biological Systems, Institute of Physics of The National Academy of Sciences of Ukraine, 46 Nauky Ave., 03038, Kyiv, Ukraine.

[3] Department of Biophysics and Medical Informatics, Educational and Scientific Centre "Institute of Biology and Medicine", Taras Shevchenko Kyiv National University, 64 Volodymyrska Str., 01601, Kyiv, Ukraine.

[4] Department of Cellular, Computational and Integrative Biology, The University of Trento, Via Sommarive 9, 38123 Povo (Trento), Italy.

[5] Institute of Organic Chemistry and Biochemistry, Czech Academy of Sciences, CZ-166 10 Prague 6, Czech Republic.

[6] Department of Physical Chemistry, Faculty of Science, Palacký University Olomouc, 17. listopadu 12, 771 46 Olomouc, Czech Republic.

*

Table 1. Main characteristics of TDC ADMET endpoints

| Category | Benchmark | Task | Metric | Train Size | Test Size | Description |
|---|---|---|---|---|---|---|
| **Absorption** | caco2_wang | Regression | MAE | 706 | 176 | Caco-2 cell effective permeability |
| | bioavailability_ma | Classification | AUROC | 475 | 119 | Oral bioavailability (F > 20%) |
| | lipophilicity_astrazeneca | Regression | MAE | 3,360 | 840 | Lipophilicity (log D7.4) |
| | solubility_aqsoldb | Regression | MAE | 7,831 | 1,958 | Aqueous solubility |
| **Distribution** | hia_hou | Classification | AUROC | 475 | 119 | Human intestinal absorption |
| | pgp_broccatelli | Classification | AUROC | 1,028 | 257 | P-glycoprotein inhibition |
| | bbb_martins | Classification | AUROC | 1,560 | 390 | Blood-brain barrier penetration |
| | ppbr_az | Regression | MAE | 1,520 | 380 | Plasma protein binding rate |
| **Metabolism** | vdss_lombardo | Regression | Spearman | 968 | 242 | Volume of distribution |
| | cyp2c9_veith | Classification | AUPRC | 10,192 | 2,548 | CYP2C9 inhibition |
| | cyp2d6_veith | Classification | AUPRC | 11,127 | 2,782 | CYP2D6 inhibition |
| | cyp3a4_veith | Classification | AUPRC | 10,164 | 2,541 | CYP3A4 inhibition |
| | cyp2c9_substrate_carbonmangels | Classification | AUPRC | 525 | 131 | CYP2C9 substrate |
| | cyp2d6_substrate_carbonmangels | Classification | AUPRC | 571 | 143 | CYP2D6 substrate |
| | cyp3a4_substrate_carbonmangels | Classification | AUROC | 534 | 134 | CYP3A4 substrate |
| **Excretion** | half_life_obach | Regression | Spearman | 534 | 134 | Terminal half-life |
| | clearance_hepatocyte_az | Regression | Spearman | 870 | 218 | Hepatocyte intrinsic clearance |
| | clearance_microsome_az | Regression | Spearman | 1,020 | 255 | Microsome intrinsic clearance |
| **Toxicity** | ld50_zhu | Regression | MAE | 5,931 | 1,483 | Acute oral toxicity LD50 |
| | herg | Classification | AUROC | 538 | 135 | hERG cardiotoxicity |
| | ames | Classification | AUROC | 6,265 | 1,566 | AMES mutagenicity |
| | dili | Classification | AUROC | 382 | 96 | Drug-induced liver injury |

Table S2. Fingerprint types - candidates for sequential forward selection

| Category | Fingerprint | Abbreviation | Size | Parameters |
|---|---|---|---|---|
| **Binary (folded)** | Extended Connectivity Fingerprint | ecfp | 1024 | radius=2 |
| | Functional Connectivity Fingerprint | fcfp | 1024 | radius=2 |
| | Avalon Fingerprint | avalon | 1024 | - |
| | RDKit Fingerprint | rdkit | 1024 | - |
| | Topological Torsion Fingerprint | topological | 1024 | - |
| | Atom Pair Fingerprint | atompair | 1024 | - |
| | Pattern Fingerprint | pattern | 1024 | - |
| | Layered Fingerprint | layered | 1024 | - |
| | SMILES Extended Connectivity Fingerprint | secfp | 1024 | - |
| **Count-based (folded)** | ECFP with counts | ecfp-count | 1024 | radius=2 |
| | FCFP with counts | fcfp-count | 1024 | radius=2 |
| | RDKit with counts | rdkit-count | 1024 | - |
| | Topological Torsion with counts | topological-count | 1024 | - |
| | Atom Pair with counts | atompair-count | 1024 | - |
| **Fixed-size** | MACCS Keys | maccs | 167 | - |
| | Extended Reduced Graph | erg | 315 | - |
| | E-State Fingerprint | estate | 79 | - |
| | RDKit 2D Descriptors (vectorized) | desc2D | 223 | - |
| | Mordred 2D Descriptors (vectorized) | mordred | 1613 | ignore_3D=True |
| | CATS2D | cats2D | 189 | - |
| | Scaffold Keys | scaffoldkeys | 42 | - |
| | SKeys | skeys | 42 | - |

#### Errors when setting up the required execution environment for the ADMETRix model.

```
Pip subprocess error:

ERROR: Ignored the following versions that require a different python version: 0.12.0 Requires-Python >=3.11; 0.12.0rc1 Requires-Python >=3.11; 0.12.0rc2 Requires-Python >=3.11; 0.12.0rc3 Requires-Python >=3.11; 0.12.0rc4 Requires-Python >=3.11; 0.12.1 Requires-Python >=3.11; 0.12.2 Requires-Python >=3.11; 0.12.3 Requires-Python >=3.11; 0.12.4 Requires-Python >=3.11; 0.12.5 Requires-Python >=3.11; 0.12.6 Requires-Python >=3.11; 0.12.7 Requires-Python >=3.11; 0.36.0 Requires-Python >=3.6,<3.10; 0.37.0 Requires-Python >=3.7,<3.10; 0.52.0 Requires-Python >=3.6,<3.9; 0.52.0rc3 Requires-Python >=3.6,<3.9; 0.53.0 Requires-Python >=3.6,<3.10; 0.53.0rc1.post1 Requires-Python >=3.6,<3.10; 0.53.0rc2 Requires-Python >=3.6,<3.10; 0.53.0rc3 Requires-Python >=3.6,<3.10; 0.53.1 Requires-Python >=3.6,<3.10; 0.54.0 Requires-Python >=3.7,<3.10; 0.54.0rc2 Requires-Python >=3.7,<3.10; 0.54.0rc3 Requires-Python >=3.7,<3.10; 0.54.1 Requires-Python >=3.7,<3.10; 0.7 Requires-Python >=3.6,<3.7; 0.8 Requires-Python >=3.6,<3.7; 1.3.3 Requires-Python >=3.11; 1.6.1.dev0 Requires-Python >=3.7,<3.9; 1.7.0 Requires-Python >=3.7,<3.9; 1.7.1 Requires-Python >=3.7,<3.9; 2.0.0 Requires-Python >=3.11; 2.0.0rc1 Requires-Python >=3.11; 2.0.1 Requires-Python >=3.11; 2.0.2 Requires-Python >=3.11; 2.0.3 Requires-Python >=3.11; 2.0.4 Requires-Python >=3.11; 2.0.5 Requires-Python >=3.11; 2.1.0 Requires-Python >=3.11; 2.1.1 Requires-Python >=3.11; 2.1.2 Requires-Python >=3.11; 2.2.0 Requires-Python >=3.11; 2.2.1 Requires-Python >=3.11; 2.3.0 Requires-Python >=3.11; 2.3.1 Requires-Python >=3.11; 2.3.2 Requires-Python >=3.11; 2.3.3 Requires-Python >=3.11; 2.3.4 Requires-Python >=3.11; 2.3.5 Requires-Python >=3.11; 2.4.0 Requires-Python >=3.11; 2.4.0rc1 Requires-Python >=3.11; 2.4.1 Requires-Python >=3.11; 3.0.0rc0 Requires-Python >=3.11; 3.0.0rc1 Requires-Python >=3.11; 3.5 Requires-Python >=3.11; 3.5rc0 Requires-Python >=3.11; 3.6 Requires-Python >=3.11; 3.6.1 Requires-Python >=3.11,!=3.14.1; 3.6rc0 Requires-Python >=3.11; 9.0.0 Requires-Python >=3.11; 9.0.0b1 Requires-Python >=3.11; 9.0.0b2 Requires-Python >=3.11; 9.0.1 Requires-Python >=3.11; 9.0.2 Requires-Python >=3.11; 9.1.0 Requires-Python >=3.11; 9.2.0 Requires-Python >=3.11; 9.3.0 Requires-Python >=3.11; 9.4.0 Requires-Python >=3.11; 9.5.0 Requires-Python >=3.11; 9.6.0 Requires-Python >=3.11; 9.7.0 Requires-Python >=3.11; 9.8.0 Requires-Python >=3.11; 9.9.0 Requires-Python >=3.11

ERROR: Could not find a version that satisfies the requirement python-graphviz==0.20.3 (from versions: none)

ERROR: No matching distribution found for python-graphviz==0.20.3

failed
```

#### Errors when setting up the required execution environment for the SimGCN model.

```
Traceback (most recent call last):
 File "/home/nazar/Desktop/TDC/SimGCN-TDC/run.py", line 5, in <module>
   from katanaHLS.data.dataset import SIPGraph
```

```
 File "/home/nazar/Desktop/TDC/SimGCN-TDC/katanaHLS/data/dataset.py", line
8, in <module>
   from torch_geometric.data import InMemoryDataset, Data
File
"/home/nazar/.pyenv/versions/3.12.3/lib/python3.12/site-packages/torch_geo
metric/__init__.py", line 7, in <module>
   import torch_geometric.data
File
"/home/nazar/.pyenv/versions/3.12.3/lib/python3.12/site-packages/torch_geo
metric/data/__init__.py", line 1, in <module>
   from .data import Data
 File
"/home/nazar/.pyenv/versions/3.12.3/lib/python3.12/site-packages/torch_geo
metric/data/data.py", line 3, in <module>
   from torch_geometric.typing import OptTensor, NodeType, EdgeType
 File
"/home/nazar/.pyenv/versions/3.12.3/lib/python3.12/site-packages/torch_geo
metric/typing.py", line 4, in <module>
   from torch_sparse import SparseTensor
ModuleNotFoundError: No module named 'torch_sparse'
 Cant install torch sparse
Collecting torch-sparse
  Using cached torch_sparse-0.6.18.tar.gz (209 kB)
  Installing build dependencies ... done
  Getting requirements to build wheel ... error
  error: subprocess-exited-with-error
 × Getting requirements to build wheel did not run successfully.
  │ exit code: 1
  ╰─> [20 lines of output]
      Traceback (most recent call last):
        File
"/home/nazar/.pyenv/versions/3.12.3/lib/python3.12/site-packages/pip/_vend
or/pyproject_hooks/_in_process/_in_process.py", line 389, in <module>
          main()
        File
"/home/nazar/.pyenv/versions/3.12.3/lib/python3.12/site-packages/pip/_vend
or/pyproject_hooks/_in_process/_in_process.py", line 373, in main
          json_out["return_val"] = hook(**hook_input["kwargs"])
                                   ^^^^^^^^^^^^^^^^^^^^^^^^^^^^
        File
"/home/nazar/.pyenv/versions/3.12.3/lib/python3.12/site-packages/pip/_vend
or/pyproject_hooks/_in_process/_in_process.py", line 143, in
get_requires_for_build_wheel
          return hook(config_settings)
                 ^^^^^^^^^^^^^^^^^^^^^
        File
"/tmp/pip-build-env-1ooew3s2/overlay/lib/python3.12/site-packages/setuptoo
ls/build_meta.py", line 331, in get_requires_for_build_wheel
          return self._get_build_requires(config_settings,
requirements=[])

^^^^^^^^^^^^^^^^^^^^^^^^^^^^^^^^^^^^^^^^^^^^^^^^^^^^^^^^^^
        File
"/tmp/pip-build-env-1ooew3s2/overlay/lib/python3.12/site-packages/setuptoo
ls/build_meta.py", line 301, in _get_build_requires
          self.run_setup()
        File
"/tmp/pip-build-env-1ooew3s2/overlay/lib/python3.12/site-packages/setuptoo
ls/build_meta.py", line 512, in run_setup
          super().run_setup(setup_script=setup_script)
```

```
        File
"/tmp/pip-build-env-1ooew3s2/overlay/lib/python3.12/site-packages/setuptoo
ls/build_meta.py", line 317, in run_setup
          exec(code, locals())
        File "<string>", line 8, in <module>
      ModuleNotFoundError: No module named 'torch'
      [end of output]
 note: This error originates from a subprocess, and is likely not a
problem with pip.
ERROR: Failed to build 'torch-sparse' when getting requirements to build
wheel

In /usr/lib/x86_64-linux-gnu run something like sudo ln -s libtiff.so.6
libtiff.so.5
doesn't work
```

#### ZairaChem runtime errors

```
(base) nazar_shevchuk_receptor_ai@admet-predict:~/zaira-chem$ conda
activate zairachem
(zairachem) nazar_shevchuk_receptor_ai@admet-predict:~/zaira-chem$
zairachem --help
2026-02-06 02:24:30.101954: I
tensorflow/core/platform/cpu_feature_guard.cc:193] This TensorFlow binary
is optimized with oneAPI Deep Neural Network Library (oneDNN) to use the
following CPU instructions in performance-critical operations:  AVX2 FMA
To enable them in other operations, rebuild TensorFlow with the
appropriate compiler flags.
2026-02-06 02:24:30.314378: E
tensorflow/stream_executor/cuda/cuda_blas.cc:2981] Unable to register
cuBLAS factory: Attempting to register factory for plugin cuBLAS when one
has already been registered
2026-02-06 02:24:31.237772: W
tensorflow/stream_executor/platform/default/dso_loader.cc:64] Could not
load dynamic library 'libnvinfer.so.7'; dlerror: libnvinfer.so.7: cannot
open shared object file: No such file or directory
2026-02-06 02:24:31.237930: W
tensorflow/stream_executor/platform/default/dso_loader.cc:64] Could not
load dynamic library 'libnvinfer_plugin.so.7'; dlerror:
libnvinfer_plugin.so.7: cannot open shared object file: No such file or
directory
2026-02-06 02:24:31.237952: W
tensorflow/compiler/tf2tensorrt/utils/py_utils.cc:38] TF-TRT Warning:
Cannot dlopen some TensorRT libraries. If you would like to use Nvidia GPU
with TensorRT, please make sure the missing libraries mentioned above are
installed properly.
Traceback (most recent call last):
  File
"/home/nazar_shevchuk_receptor_ai/miniconda3/envs/zairachem/bin/zairachem"
, line 3, in <module>
    from zairachem.cli import cli
  File
"/home/nazar_shevchuk_receptor_ai/zaira-chem/zairachem/cli/__init__.py",
line 1, in <module>
```

```
    from .create_cli import create_cli
  File
"/home/nazar_shevchuk_receptor_ai/zaira-chem/zairachem/cli/create_cli.py",
line 1, in <module>
    from .cmd import Command
  File "/home/nazar_shevchuk_receptor_ai/zaira-chem/zairachem/cli/cmd.py",
line 4, in <module>
    from .commands.estimate import estimate_cmd
  File
"/home/nazar_shevchuk_receptor_ai/zaira-chem/zairachem/cli/commands/estima
te.py", line 7, in <module>
    from ...estimators.pipe import EstimatorPipeline
  File
"/home/nazar_shevchuk_receptor_ai/zaira-chem/zairachem/estimators/pipe.py"
, line 10, in <module>
    from .from_individual_full_descriptors_tabpfn.pipe import (
  File
"/home/nazar_shevchuk_receptor_ai/zaira-chem/zairachem/estimators/from_ind
ividual_full_descriptors_tabpfn/pipe.py", line 1, in <module>
    from .estimate import Estimator
  File
"/home/nazar_shevchuk_receptor_ai/zaira-chem/zairachem/estimators/from_ind
ividual_full_descriptors_tabpfn/estimate.py", line 12, in <module>
    from ...automl.binarytabpfn import TabPFNBinaryClassifier
  File
"/home/nazar_shevchuk_receptor_ai/zaira-chem/zairachem/automl/binarytabpfn
.py", line 5, in <module>
    from tabpfn import TabPFNClassifier
  File
"/home/nazar_shevchuk_receptor_ai/miniconda3/envs/zairachem/lib/python3.10
/site-packages/tabpfn/__init__.py", line 1, in <module>
    from tabpfn.scripts.transformer_prediction_interface import
TabPFNClassifier
  File
"/home/nazar_shevchuk_receptor_ai/miniconda3/envs/zairachem/lib/python3.10
/site-packages/tabpfn/scripts/transformer_prediction_interface.py", line
19, in <module>
    from tabpfn.scripts.model_builder import load_model,
load_model_only_inference
  File
"/home/nazar_shevchuk_receptor_ai/miniconda3/envs/zairachem/lib/python3.10
/site-packages/tabpfn/scripts/model_builder.py", line 4, in <module>
    from tabpfn.transformer import TransformerModel
  File
"/home/nazar_shevchuk_receptor_ai/miniconda3/envs/zairachem/lib/python3.10
/site-packages/tabpfn/transformer.py", line 9, in <module>
    from tabpfn.layer import TransformerEncoderLayer, _get_activation_fn
  File
"/home/nazar_shevchuk_receptor_ai/miniconda3/envs/zairachem/lib/python3.10
/site-packages/tabpfn/layer.py", line 5, in <module>
    from torch.nn.modules.transformer import _get_activation_fn, Module,
Tensor, Optional, MultiheadAttention, Linear, Dropout, LayerNorm
```

```
ImportError: cannot import name 'Optional' from
'torch.nn.modules.transformer'
(/home/nazar_shevchuk_receptor_ai/miniconda3/envs/zairachem/lib/python3.10
/site-packages/torch/nn/modules/transformer.py)
```

Table S3. Metrics and ranks of in-house models on different TDC ADMET endpoints

| benchmark | metric name | baseline metric | baseline rank | sfs metric | sfs rank | hpo metric | hpo rank | total | features | best_params |
|---|---|---|---|---|---|---|---|---|---|---|
| caco2_wang | MAE | 0.3067 | 9 | 0.2889 | 7 | 0.2875 | 7 | 24 | mol2vec, filtered_descs, topological-count, maccs | n_estimators 200<br>learning_rate 0.1<br>max_depth -1<br>num_leaves 63 |
| bioavailability_ma | AUROC | 0.6784 | 21 | 0.6744 | 21 | 0.6744 | 21 | 20 | mol2vec, topological-count, rdkit-count, topological, layered | n_estimators 100<br>learning_rate 0.1<br>max_depth -1<br>num_leaves 31 |
| lipophilicity_astrazeneca | MAE | 0.5701 | 13 | 0.5225 | 7 | 0.5108 | 5 | 21 | mol2vec, filtered_descs, avalon-count, ecfp-count, estate | n_estimators 800<br>learning_rate 0.05<br>max_depth 5<br>num_leaves 15 |
| solubility_aqsoldb | MAE | 0.813 | 7 | 0.7932 | 7 | 0.7808 | 5 | 18 | mol2vec, mordred, avalon-count, desc2D, topological-count | n_estimators 400<br>learning_rate 0.03<br>max_depth -1<br>num_leaves 63 |
| hia_hou | AUROC | 0.9901 | 2 | 0.9938 | 1 | 0.9938 | 1 | 20 | mol2vec, mordred, ecfp-count, maccs, pattern | |

| | | | | | | | | | | |
|---|---|---|---|---|---|---|---|---|---|---|
| pgp_broccatelli | AUROC | 0.9123 | 8 | 0.8968 | 10 | 0.902 | 10 | 16 | mol2vec, filtered_descs, erg, atompair-count, estate | n_estimators 800<br>learning_rate 0.1<br>max_depth 9<br>num_leaves 127 |
| bbb_martins | AUROC | 0.9199 | 8 | 0.9225 | 2 | 0.8884 | 17 | 26 | mol2vec, erg, pattern | n_estimators 100<br>learning_rate 0.01<br>max_depth 9<br>num_leaves 31 |
| ppbr_az | MAE | 8.4496 | 9 | 8.4795 | 10 | 7.9457 | 7 | 20 | mol2vec, mordred, pattern, desc2D | n_estimators 400<br>learning_rate 0.05<br>max_depth 5<br>num_leaves 15 |
| vdss_lombardo | Spearman | 0.3356 | 17 | 0.3368 | 17 | 0.4949 | 11 | 19 | mol2vec, desc2D, rdkit-count, topological, mordred | n_estimators 100<br>learning_rate 0.01<br>max_depth -1<br>num_leaves 63 |
| cyp2c9_veith | AUPRC | 0.7613 | 9 | 0.7883 | 6 | 0.7922 | 5 | 20 | mol2vec, mordred, rdkit-count, avalon-count, scaffoldkeys | n_estimators 800<br>learning_rate 0.1<br>max_depth 9<br>num_leaves 63 |
| cyp2d6_veith | AUPRC | 0.7002 | 7 | 0.7069 | 7 | 0.7178 | 7 | 19 | mol2vec, filtered_descs, ecfp-count, topological, avalon-count | n_estimators 800<br>learning_rate 0.05<br>max_depth -1<br>num_leaves 63 |
| cyp3a4_veith | AUPRC | 0.8511 | 11 | 0.8683 | 9 | 0.8748 | 9 | 19 | mol2vec, avalon-count, rdkit, cats2D, scaffoldkeys | n_estimators 800<br>learning_rate 0.05<br>max_depth 3<br>num_leaves 15 |

| | | | | | | | | | | |
|---|---|---|---|---|---|---|---|---|---|---|
| cyp2c9_substrate_carbonmangels | AUPRC | 0.3363 | 20 | 0.3459 | 19 | 0.3459 | 19 | 20 | mol2vec, filtered_descs | n_estimators 100<br>learning_rate 0.1<br>max_depth -1<br>num_leaves 31 |
| cyp2d6_substrate_carbonmangels | AUPRC | 0.657 | 12 | 0.6842 | 10 | 0.6842 | 10 | 18 | mol2vec, rdkit-count, maccs, desc2D, ecfp | n_estimators 100<br>learning_rate 0.1<br>max_depth -1<br>num_leaves 31 |
| cyp3a4_substrate_carbonmangels | AUROC | 0.607 | 14 | 0.6384 | 9 | 0.6032 | 15 | 19 | mol2vec, mordred, pattern | n_estimators 200<br>learning_rate 0.01<br>max_depth 3<br>num_leaves 31 |
| half_life_obach | Spearman | 0.3173 | 11 | 0.2718 | 12 | 0.3927 | 9 | 20 | mol2vec, rdkit-count, avalon, atompair-count | n_estimators 200<br>learning_rate 0.01<br>max_depth -1<br>num_leaves 127 |
| clearance_hepatocyte_az | Spearman | 0.3259 | 16 | 0.3373 | 16 | 0.3353 | 16 | 18 | mol2vec, filtered_descs, scaffoldkeys, topological-count | n_estimators 100<br>learning_rate 0.05<br>max_depth -1<br>num_leaves 31 |
| clearance_microsome_az | Spearman | 0.5432 | 14 | 0.5542 | 14 | 0.5512 | 15 | 20 | mol2vec, filtered_descs, rdkit-count, layered, topological | n_estimators 400<br>learning_rate 0.1<br>max_depth -1<br>num_leaves 31 |
| ld50_zhu | MAE | 0.6212 | 6 | 0.61 | 6 | 0.5866 | 4 | 22 | mol2vec, mordred, avalon, rdkit, atompair-count | n_estimators 400<br>learning_rate 0.05<br>max_depth 9<br>num_leaves 31 |

| | | | | | | | | | | |
|---|---|---|---|---|---|---|---|---|---|---|
| herg | AUROC | 0.815 | 11 | 0.8032 | 11 | 0.8032 | 11 | 20 | mol2vec, filtered_descs, maccs, erg, cats2D | n_estimators 100<br>learning_rate 0.1<br>max_depth -1<br>num_leaves 31 |
| ames | AUROC | 0.8255 | 13 | 0.8422 | 10 | 0.7942 | 17 | 20 | mol2vec, pattern, layered, cats2D, rdkit | n_estimators 100<br>learning_rate 0.01<br>max_depth -1<br>num_leaves 15 |
| dili | AUROC | 0.8787 | 13 | 0.86 | 16 | 0.86 | 16 | 20 | mol2vec, avalon, atompair, pattern | n_estimators 100<br>learning_rate 0.1<br>max_depth -1<br>num_leaves 31 |

Table S4. Metrics and ranks of deliberately overfitted in-house models on different TDC ADMET endpoints

| **benchmark** | **metric name** | **baseline metric** | **baseline rank** | **sfs metric** | **sfs rank** | **hpo metric** | **hpo rank** | **total** | **features** | **best_params** |
|---|---|---|---|---|---|---|---|---|---|---|
| caco2_wang | MAE | 0.3067 | 9 | 0.2694 | 2 | 0.2694 | 2 | 24 | mol2vec, desc2D, erg, filtered_descs | n_estimators 100<br>learning_rate 0.1 |
| bioavailability_ma | AUROC | 0.6784 | 21 | 0.7147 | 21 | 0.7147 | 21 | 20 | mol2vec, topological, atompair, rdkit-count | n_estimators 100<br>learning_rate 0.1 |
| lipophilicity_astrazeneca | MAE | 0.5701 | 13 | 0.5345 | 8 | 0.5112 | 5 | 21 | mol2vec, filtered_descs, avalon-count, layered, atompair-count | n_estimators 300<br>learning_rate 0.1 |

| | | | | | | | | | | |
|---|---|---|---|---|---|---|---|---|---|---|
| solubility_aqsoldb | MAE | 0.813 | 7 | 0.7883 | 5 | 0.7735 | 3 | 18 | mol2vec, filtered_descs, mordred | n_estimators 300<br>learning_rate 0.1 |
| hia_hou | AUROC | 0.9901 | 2 | 0.9951 | 1 | 0.9951 | 1 | 20 | mol2vec, erg, maccs, pattern, mordred | |
| pgp_broccatelli | AUROC | 0.9123 | 8 | 0.9188 | 7 | 0.9202 | 7 | 16 | mol2vec, mordred, rdkit-count, cats2D | n_estimators 200<br>learning_rate 0.2 |
| bbb_martins | AUROC | 0.9199 | 8 | 0.9376 | 1 | 0.9376 | 1 | 26 | mol2vec, atompair-count, cats2D, atompair | |
| ppbr_az | MAE | 8.4496 | 9 | 7.779 | 5 | 7.9893 | 7 | 20 | mol2vec, filtered_descs, rdkit | n_estimators 300<br>learning_rate 0.05 |
| vdss_lombardo | Spearman | 0.3356 | 17 | 0.4909 | 13 | 0.4664 | 15 | 19 | mol2vec, avalon-count, filtered_descs, desc2D | n_estimators 100,<br>learning_rate 0.1 |
| cyp2c9_veith | AUPRC | 0.7613 | 9 | 0.792 | 5 | 0.792 | 5 | 20 | mol2vec, mordred, avalon-count, layered, secfp | n_estimators 100<br>learning_rate 0.1 |
| cyp2d6_veith | AUPRC | 0.7002 | 7 | 0.7197 | 5 | 0.7085 | 7 | 19 | mol2vec, filtered_descs, atompair-count | n_estimators 200<br>learning_rate 0.05 |
| cyp3a4_veith | AUPRC | 0.8511 | 11 | 0.8733 | 9 | 0.8826 | 5 | 19 | mol2vec, filtered_descs, ecfp-count, layered, pattern | n_estimators 300<br>learning_rate 0.2 |
| cyp2c9_substrate_carbonmangels | AUPRC | 0.3363 | 20 | 0.3566 | 18 | 0.3566 | 18 | 20 | mol2vec, mordred | n_estimators 100<br>learning_rate 0.1 |

| | | | | | | | | | | |
|---|---|---|---|---|---|---|---|---|---|---|
| cyp2d6_substrate_carbonmangels | AUPRC | 0.657 | 12 | 0.7566 | 1 | 0.7566 | 1 | 18 | mol2vec, desc2D, secfp, avalon-count | |
| cyp3a4_substrate_carbonmangels | AUROC | 0.607 | 14 | 0.6658 | 2 | 0.6658 | 2 | 19 | mol2vec, maccs, scaffoldkeys | n_estimators 100<br>learning_rate 0.1 |
| half_life_obach | Spearman | 0.3173 | 11 | 0.4053 | 9 | 0.4089 | 9 | 20 | mol2vec, mordred, topological-count, rdkit | n_estimators 100<br>learning_rate 0.05 |
| clearance_hepatocyte_az | Spearman | 0.3259 | 16 | 0.4413 | 6 | 0.4413 | 6 | 18 | mol2vec, secfp, ecfp, atompair | n_estimators 100<br>learning_rate 0.1 |
| clearance_microsome_az | Spearman | 0.5432 | 14 | 0.6314 | 1 | 0.6314 | 1 | 20 | mol2vec, avalon-count, desc2D, atompair | |
| ld50_zhu | MAE | 0.6212 | 6 | 0.5964 | 5 | 0.5766 | 3 | 22 | mol2vec, mordred, atompair-count, avalon, topological-count | n_estimators 300<br>learning_rate 0.1 |
| herg | AUROC | 0.815 | 11 | 0.8495 | 6 | 0.8495 | 6 | 20 | mol2vec, desc2D, rdkit-count, layered, secfp | n_estimators 100<br>learning_rate 0.1 |
| ames | AUROC | 0.8255 | 13 | 0.8652 | 5 | 0.8723 | 1 | 20 | mol2vec, mordred, rdkit, avalon-count, topological | n_estimators 300<br>learning_rate 0.1 |
| dili | AUROC | 0.8787 | 13 | 0.9422 | 2 | 0.9391 | 2 | 20 | mol2vec, mordred, filtered_descs, scaffoldkeys, secfp | n_estimators 200<br>learning_rate 0.05 |
